# Mechanistic and experimental models of cell migration reveal the importance of intercellular interactions in cell invasion

**DOI:** 10.1101/391557

**Authors:** Oleksii M. Matsiaka, Ruth E. Baker, Esha T. Shah, Matthew J. Simpson

## Abstract

Moving fronts of cells are essential for development, repair and disease progression. Therefore, understanding and quantifying the details of the mechanisms that drive the movement of cell fronts is of wide interest. Quantitatively identifying the role of intercellular interactions, and in particular the role of cell pushing, remains an open question. Indeed, perhaps the most common continuum mathematical idealization of moving cell fronts is to treat the population of cells, either implicitly or explicitly, as a population of point particles undergoing a random walk that neglects intercellular interactions. In this work, we report a combined experimental-modelling approach showing that intercellular interactions contribute significantly to the spatial spreading of a population of cells. We use a novel experimental data set with PC-3 prostate cancer cells that have been pretreated with Mitomycin-C to suppress proliferation. This allows us to experimentally separate the effects of cell migration from cell proliferation, thereby enabling us to focus on the migration process in detail as the population of cells recolonizes an initially-vacant region in a series of two-dimensional experiments. We quantitatively model the experiments using a stochastic modelling framework, based on Langevin dynamics, which explicitly incorporates random motility and various intercellular forces including: (i) long range attraction (adhesion); and (ii) finite size effects that drive short range repulsion (pushing). Quantitatively comparing the ability of this model to describe the experimentally observed population-level behaviour provides us with quantitative insight into the roles of random motility and intercellular interactions. To quantify the mechanisms at play, we calibrate the stochastic model to match experimental cell density profiles to obtain estimates of cell diffusivity, *D*, and the amplitude of intercellular forces, *f*_0_. Our analysis shows that taking a standard modelling approach which ignores intercellular forces provides a poor match to the experimental data whereas incorporating intercellular forces, including short-range pushing and longer range attraction, leads to a faithful representation of the experimental observations. These results demonstrate a significant role for intercellular interactions in cell invasion.

**Author summary:** Moving cell fronts are routinely observed in various physiological processes, such as wound healing, malignant invasion and embryonic morphogenesis. We explore the effects of a previously overlooked mechanism that contributes to population-level front movement: pushing. Our framework is flexible and incorporates range of reasonable biological phenomena, such as random motility, cell-to-cell adhesion, and pushing. We find that neglecting finite size effects and intercellular forces, such as cell pushing, reduces our ability to mimic and predict our experimental observations.

## Introduction

Moving cell fronts occur during many physiological processes, such as wound healing, developmental morphogenesis, and malignant invasion [1, 2, 3, 4, 5, 6]. Typically, cell fronts are observed as advancing, sharp boundaries between densely occupied and vacant regions, or as a moving interface between two distinct populations of cells. An example of the first scenario is wound healing, where populations of cells close and recolonize an initially vacant space [7, 8], as shown in Fig 1. An advancing interface between two populations of cells is often associated with malignant invasion into surrounding tissues [9, 10, 11]. Thus, improving our understanding of how cell populations spread can provide important, clinically-relevant information about the nature of moving cell fronts. Historically, moving cell fronts have been studied, both *in vitro* and *in vivo*, to provide both qualitative and quantitative information about the mechanisms that drive front movement. We note that quantifying the precise contributions of various cellular-level mechanisms that lead to population-level front behavior is a nontrivial task that requires the integration of many different types of experimental data [12]. Often it is assumed that the movement of advancing cell fronts is driven by combined effects of undirected cell migration and carrying capacity-limited cell proliferation [13, 12]. At present, a fundamental question, which remains largely overlooked in the mathematical biology literature, is – *what is the role of cell-to-cell interactions and how does short range cell pushing influence population-level front movement?*

**Figure 1:**
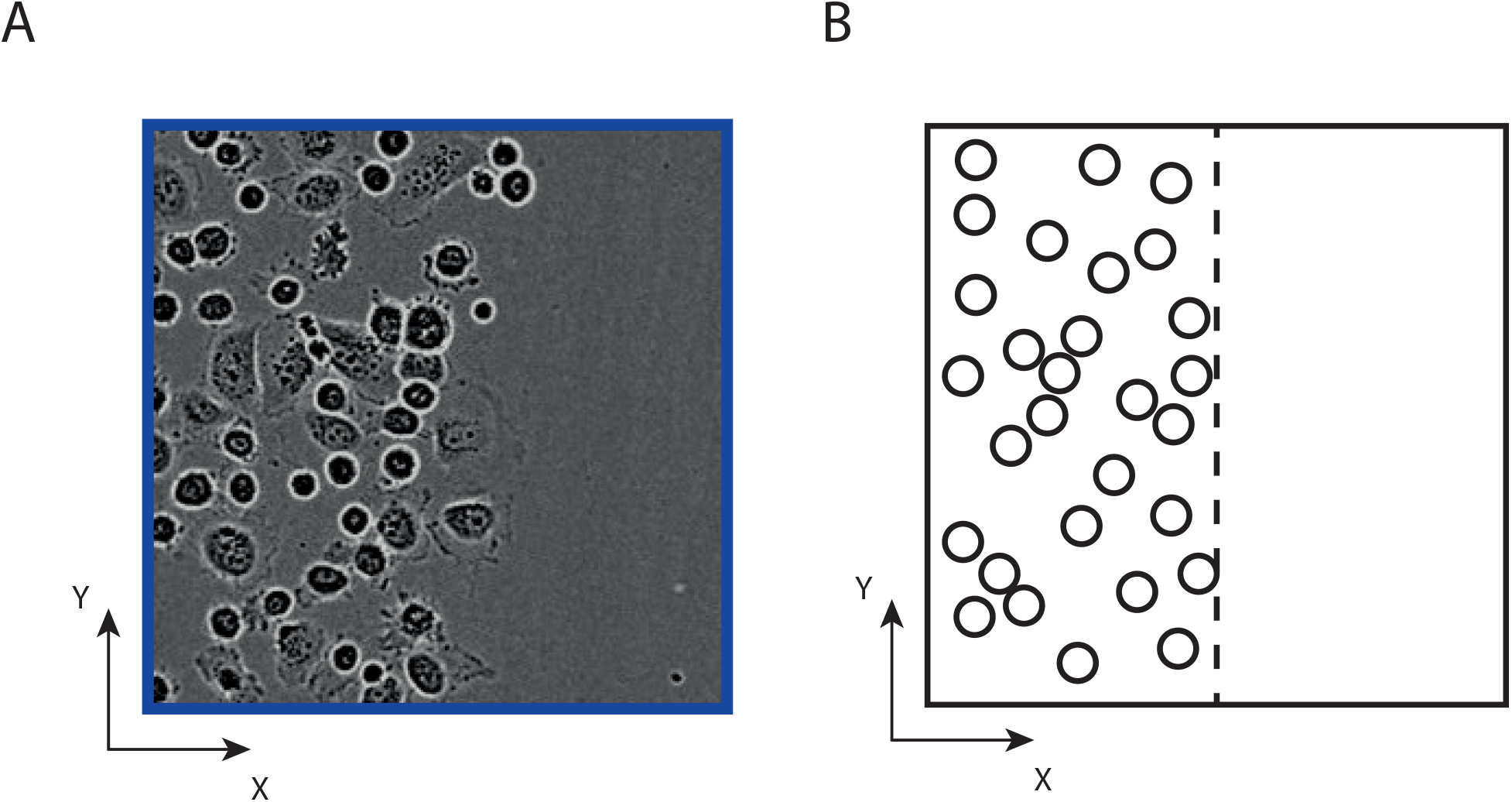
What drives the movement of cell fronts? A: Experimental image showing the leading edge of a moving front of PC-3 prostate cancer cells. This front is moving in the positive *x* direction. B: Schematic of A showing the position of the front (vertical dashed line) with the front moving in the positive *x* direction. The location of the region in A is the blue rectangle superimposed in Fig 2(A).

Cell motility is a complicated process involving both the interplay and competition between various individual-level mechanisms [14, 15, 16, ?]. One of the most well-studied individual-level cell motility mechanisms is lamellipodial cell migration where cells undergo undirected movement due to myosin-powered contractions of the actin network under the cytoplasmic membrane [17]. Since this process is observed in many cell types, it remains prevalent in many mathematical modelling frameworks. As such, the assumption that cells undergo Brownian motion is often invoked and cells are represented, either implicitly or explicitly, as non-interacting point particles that move according to a white Gaussian process [18]. While this approach is appealing due to its simplicity, it neglects the effects of intercellular interactions, such as adhesion and finite size (crowding) effects. The neglect of cell-cell adhesion can be problematic because it is known that mesenchymal cell types, such as keratinocytes, can be strongly affected by cell-cell adhesion during wound healing [19]. Furthermore, adherent cells can form clusters that exhibit qualitatively different behaviour from isolated cells [20, 21, 22]. Arguably, some of the most striking examples of front-like spreading of a cell population occur during embryonic development, such as neural crest cell invasion in the developing gut tissues, which is though to arise as a consequence of combined undirected Brownian cell motility and carrying capacity-limited cell proliferation [13, 23]. However, previous investigations have made the point that short range cell pushing can also play a role in driving the movement of cell fronts in confined environments, such as living tissues [24]. Henceforth, we hypothesize that cells in a confined space may generate population pressure, driven by finite size effects and local repulsion, which can stimulate spatial expansion of the population.

Perhaps the most popular mathematical framework for modelling the movement of cell fronts involves using reaction-diffusion equations [25, 26, 27, 28, 29], including the Fisher-Kolmogorov equation, and generalisations thereof. Although classical reaction-diffusion equations are routinely used to describe the movement of cell fronts, there are a few notable disadvantages of this framework, namely: (i) classical reaction-diffusion models based on linear diffusion do not include any cell-to-cell interaction forces; and (ii) classical reaction-diffusion models do not explicitly model individual cells within the population. Previously, the lack of individual-level experimental data meant that classical reaction-diffusion models might have been an acceptable way to conceptualize and simulate collective cell behaviour. However, with the increasing availability of individual-level information it is becoming increasingly important to develop mathematical models that provide both population-level information and individual-level information.

In this work we quantitatively examine the roles of random motility and intercellular interactions, including both long range adhesion and short range pushing, in a canonical experiment describing the movement of cell fronts on a two-dimensional substrate. Previously, cell pushing has been incorporated into lattice-based models of cell motility where agents move on a spatial domain that is represented as a regular lattice [30, 31]. However, these previous studies about the role of cell pushing are primarily theoretical studies that do not consider calibrating models to quantitatively match any experimental data. Our current work is the first attempt to incorporate intercellular interactions, including both long range adhesion and short range cell pushing, into a more realistic spatially continuous off-lattice discrete model of cell migration. Importantly, we directly apply the model to quantitatively mimic a novel experimental data set. In our experiments we isolate the role of cell migration from the effects of cell proliferation by working with a population of cells that is pre-treated with a chemotherapy drug to suppress proliferation. This is a critical feature of our work because it is well-known that carrying capacity-limited cell proliferation tends to dominate and mask the role of cell migration, and that it can be difficult to distinguish between different cell migration mechanisms in the presence of carrying capacity-limited cell proliferation [32]. Fig 2 illustrates the IncuCyte ZOOM™ scratch assay experimental protocol that we use in this work, at *t* = 0 h and *t* = 48 h. The IncuCyte ZOOM™ experimental system is an automatic live cell imaging technology which has two important advantages over classical scratch assays: (i) scratches are uniform and reproducible; and (ii) images are automatically recorded without interrupting an experiment [33]. As previously mentioned, we work with cells that are pretreated with the chemotherapy drug Mytomicin-C to block DNA replication and, consequently, suppress proliferation [34]. This means that the number of cells present in the experimental field of view over the duration of the experiment remains approximately constant. However, a side effect of Mytomicin-C pretreatment is that individual cells increase in size during the experiment as the cells prepare to proceed through the cell cycle but are unable to divide [35]. This dynamic increase in cell size, which is typically neglected in previous modelling studies [35], can significantly influence intercellular interactions during the experiment and so we take great care to incorporate these effects into our mathematical model. We find that our approach for incorporating dynamic cell size effects in the model is justified by also working with a simpler model that neglects dynamic changes in cell size and we find that the simpler, standard model leads to a poor match with the experimental data.

**Figure 2:**
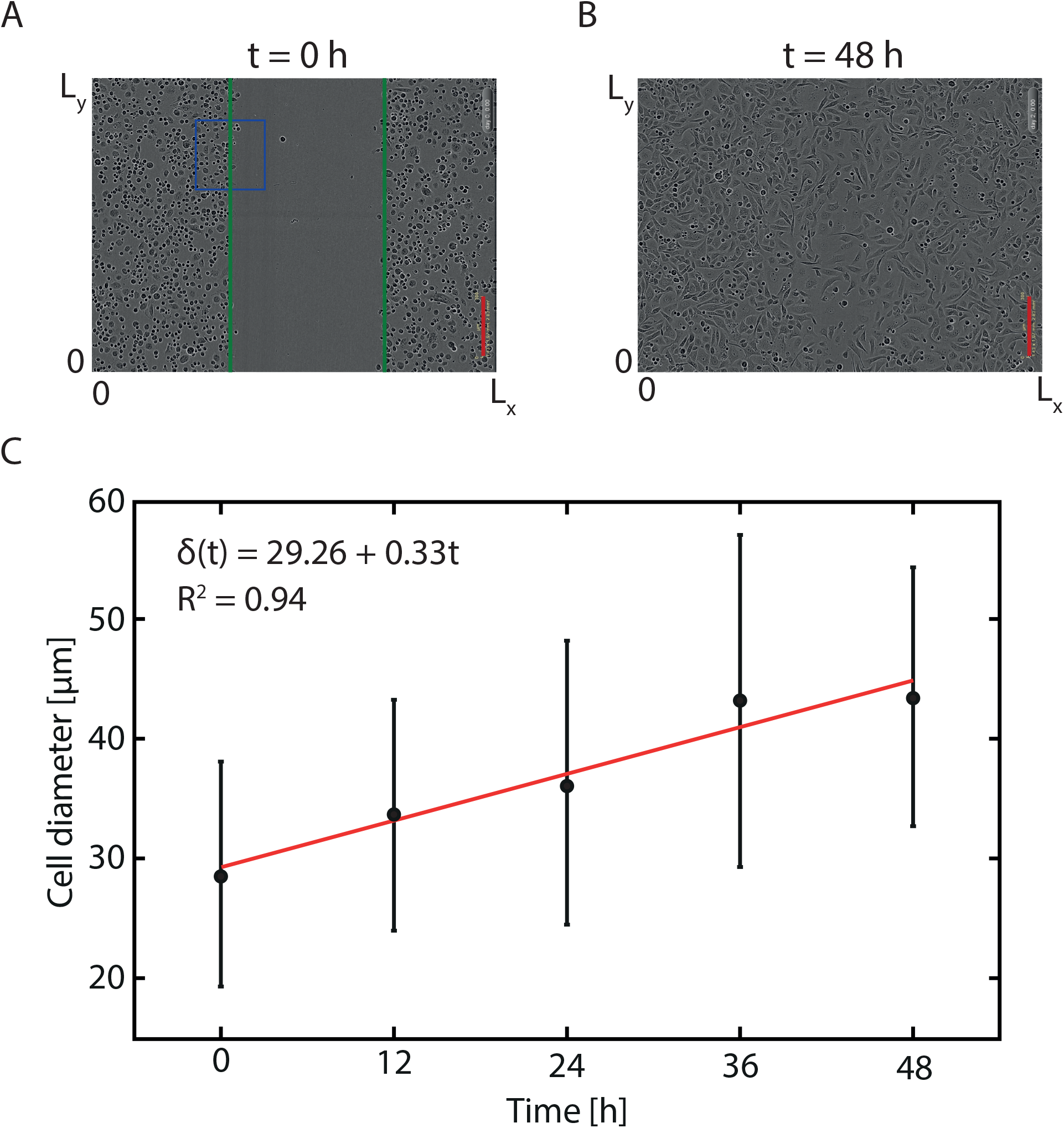
Experimental data. A-B: Images from an IncuCyte ZOOM™ assay with Mitomycin-C pretreated PC-3 prostate cancer cells. The scale bar in each image corresponds to 300 μm. The green solid lines show the initial position of the two opposingly directed cell fronts. The blue rectangle denotes the location of the subregion highlighted in Fig 1(A). C: Cell diameter data as a function of time, *δ*(*t*), from a sample of 30 randomly chosen cells at each time point. Black dots indicate the sample mean and the error bars denote the sample standard deviation about the sample mean. Red solid line represents the best-fit linear approximation, *δ*(*t*) = 29.26 + 0.33t. *R*^2^ is the adjusted coefficient of determination measuring the goodness of fit.

This work is structured as follows. We begin by describing the IncuCyte ZOOM™ experimental protocol, the experimental data, and the procedure we use to process the experimental images. We then introduce the discrete mathematical model which accounts for random motility and intercellular interactions including short range pushing and longer range attraction, as well as incorporating a mechanism for describing dynamic cell size changes. We refer to this model as *Model I* since it incorporates all four mechanisms that are thought to be relevant to the experimental system. To quantitatively explore the significance of these various cell-level mechanisms we systematically repeat the model calibration process for a range of simpler, more commonly used models. These simplified models account for: (i) random motility and intercellular forces (*Model II*); (ii) intercellular forces only (*Model III*); and (iii) random motility only (*Model IV*). We discuss the performance of each model when applied to the IncuCyte ZOOM™ data in the Results and Discussion section. Finally, in the Conclusions we summarize our findings and discuss alternative applications and extensions of our modelling framework.

## Materials and methods

### IncuCyte ZOOM™ experimental data

Monolayer scratch assays are performed using the IncuCyte ZOOM™ system (Essen BioScience, MI USA [33]) as shown in Fig 2(A-B). This technology automatically captures images of cell cultures without disrupting the fragile micro-environment. All experiments are performed using the PC-3 prostate cancer cell line [36] from the American Type Culture Collection (ATCC, Manassas, USA). The cell culture is prepared in RPMI 1640 medium (Life Technologies, Australia) in 10 % foetal calf serum (Sigma-Aldrich, Australia), with 100 U/mL penicillin, 100 g/mL streptomycin (Life Technologies), in plastic flasks (Corning Life Sciences, Asia Pacific) that are maintained in 5 % CO_2_ and 95 % air in a Panasonic incubator (VWR International) at 37 C. Cells are regularly screened for Mycoplasma (Nested PCR using primers from Sigma-Aldrich). Cells are removed from the flask using TrypLE™ (Life Technologies) in phosphate buffered saline, resuspended in medium and seeded at a density of 20,000 cells per well in 96-well ImageLock plates (Essen BioScience) as shown in Fig 3). The diameter of each individual well is 9000 μm.

**Figure 3:**
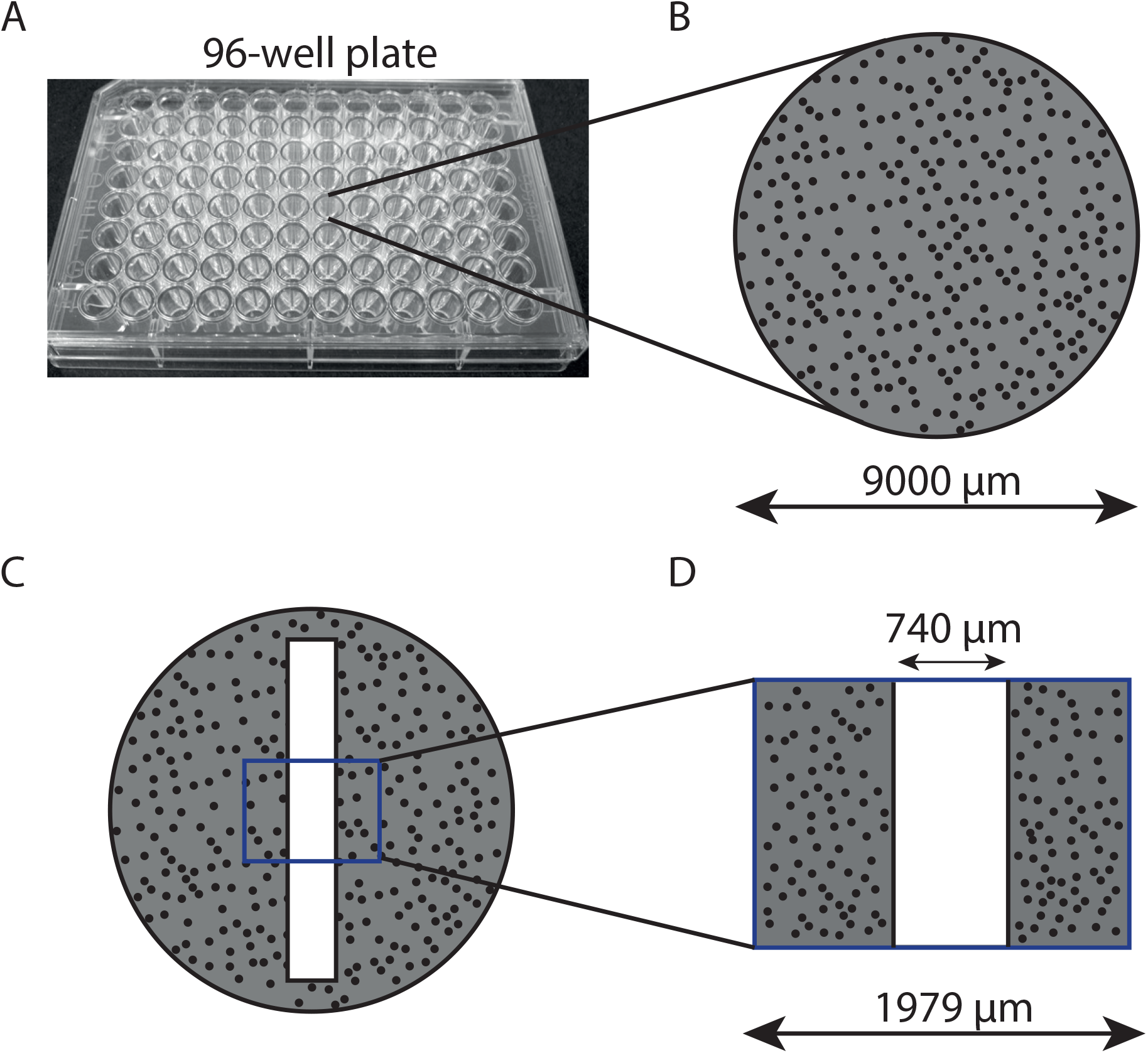
Experimental geometry. A: Image of a 96-well culture plate. The diameter of each well is 9000 μm. B: Schematic demonstrating the monolayer of cells (black dots) with approximately constant density. C: Schematic showing an artificial wound (white) in the monolayer of cells. D: Field of view of the experimental images showing that the field of view is much smaller than the extent of the well in the 96-well plate.

After seeding, cells are grown overnight to form a spatially uniform monolayer. Mitomycin-C is added at a concentration of 10 g/mL for two hours before a scratch is made in the monolayer of cells [34, 37]. A WoundMaker™ (Essen BioScience) is used to create identical scratches in the uniformly distributed populations. After creating the scratch, the medium is aspirated and wells are washed twice with fresh medium. After these washes, 100 L of fresh medium is added to each well. Once fresh medium is added, the plate is placed into the IncuCyte ZOOM™ apparatus and images of are captured every two hours for a total duration of 48 hours. In total, these experiments are performed in eight of the 96 wells on the 96-well plate. After a preliminary visual inspection of the resulting eight experimental images, we selected four typical wells for analysis. Throughout this work we will refer to these four identically prepared experiments as *Experiment A*, *Experiment B*, *Experiment C* and *Experiment D*.

Comparing images across the 48 hour duration of the experiment, summarized in Fig 2(A-B), we see that each opposingly-directed cell front moves together to close the initially-vacant region created by the scratch. By the end of the experiment we see that the Mitomycin-C pretreated cells have approximately doubled in size [35]. To quantify this increase in cell size we randomly choose 30 cells from the experimental images at *t* = 0, 12, 24, 36 and 48 h and use these cells to estimate the average diameter as a function of time, *δ*(*t*). To do this we estimate the area on the image occupied by each particular cell, and then convert this estimate of area into an equivalent diameter, 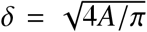, where *A* is the area estimate. With 30 estimates of the diameter at *t* = 0, 12, 24, 36 and 48 h, we compute the sample mean and sample standard deviation at each time point and plot the data in Fig 2(C). Visually we see that the average diameter appears to increase approximately linearly with time, and so we fit a linear model to the data. The cell diameter data and the linear model are shown in Fig 2(C). Note that had our experiments been performed over a longer period of time it would be better to use a different model, such as the logistic growth function, to model the temporal cell size dynamics.

### Image analysis

The experimental data used in our work is provided in the form of images from the IncuCyte ZOOM™ system. An example of a raw experimental image and a detailed description of the procedure we use to extract density information from that image are summarized in Fig 4. Here, the size of the field of view is (*L_x_ × L_y_*) = (1979 *μ*m × 1439 *μ*m), as shown in Fig 4(A). Throughout this work we use data from four identically-prepared experimental replicates of the scratch assay. Experimental images at the beginning of the experiment for each of the four replicates are shown in Fig 5. These four experimental replicates are performed simultaneously under identical experimental conditions. For completeness, the time series of experimental images for all four identically prepared experiments is given in the Supporting Information (S1 Fig). This includes images recorded at five equally-spaced time points for each of the four identically prepared experiments, giving a total of 20 experimental images.

**Figure 4:**
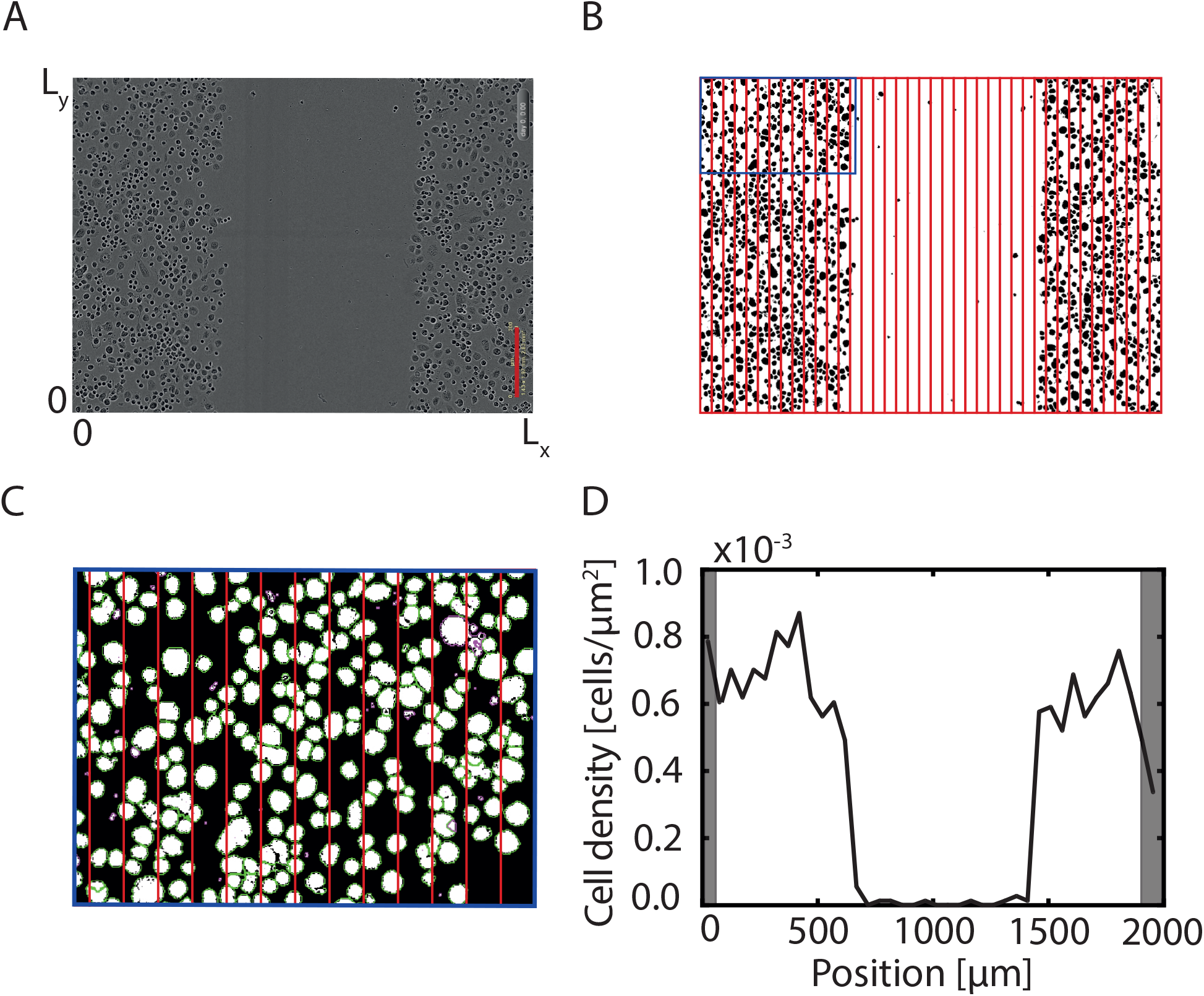
Processing of raw experimental images from the IncuCyte ZOOM™ scratch assay. A: Raw experimental image showing the field of view, of dimension (*L_x_ × L_y_*) = (1979 *μ*m × 1439 *μ*m). The scale bar corresponds to 300 μm. B: Binary image obtained after segmenting the raw image with Ilastik. C: Zoomed-in image showing the region contained within the blue rectangle in B after processing with CellProfiler. Feint green outlines denote individual objects that are identified as cells. D: The one-dimensional cell density profile is estimated by counting the number of objects per equally-spaced column. Shaded regions in D show boundary regions that are neglected owing to the presence of scale bar and time label that are automatically superimposed on the IncuCyte ZOOM™ images.

**Figure 5:**
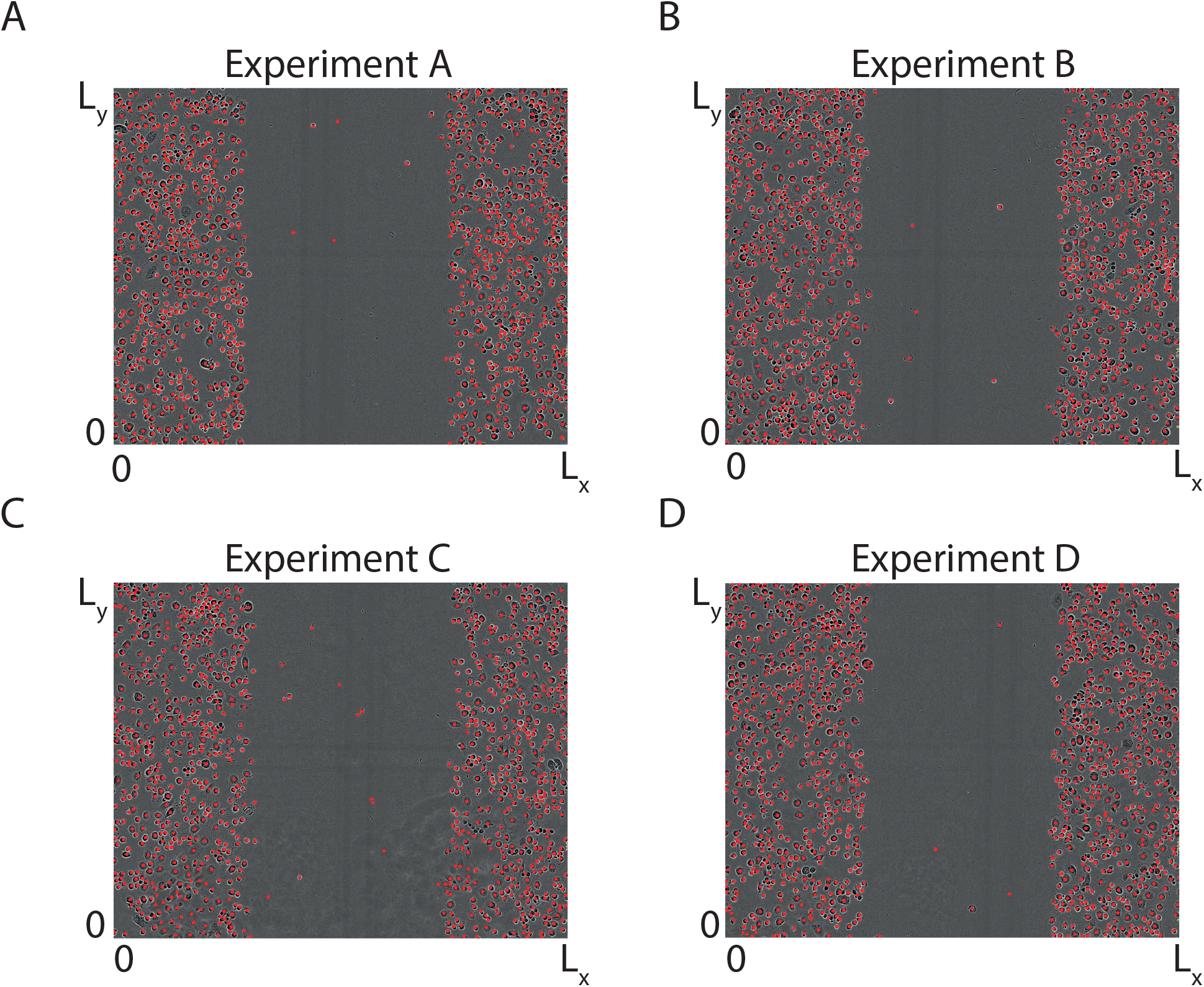
Experimental images of four identically prepared IncuCyte ZOOM™ scratch assays just after the scratch has been made, *t* = 0 h. A-D: In each experimental replicate the positions of individual cells are extracted using CellProfiler and highlighted with a red dot.

To process each experimental image we first separate the background of the image from the cells using Ilastik [38]. Ilastik is a machine learning tool that enables automatic object identification, and allows us to separate the cells in each image from the background. An example of a grey scale segmented image is shown in Fig 4(B). A visual comparison of the raw image in Fig 4(A) with the segmented image in Fig 4(B) confirms that the identification of cells from the background in the image is accurate. Since the density of cells in the original images is independent of the vertical coordinate, as shown in Fig 5, we divide each image into 40 equally-spaced columns as shown in Fig 4(B). Working with 40 columns means that the width of each column is 49.5 *μ*m. We then use CellProfiler to automatically estimate the number of cells per column [39]. With this data we divide the number of cells per column by the area of each column to give an estimate of the cell density across the horizontal coordinate of the experimental images.

A typical column-averaged cell density profile, shown in Fig 4(D), summarizes the spatial variations in density as a function of the horizontal coordinate for the particular arrangement of cells in Fig 4(A). We employ this technique to extract density profiles at each time point, *t* = 0, 12, 24, 36, and 48 h. This process is repeated four times for each of the four experimental replicates. The data presented in Fig 4(D) shows the average cell density that we associate with the centre of each column. This particular density profile is relatively noisy because it is associated with the single image in Fig 4(A). Before we proceed, we discard density data from the two right-most columns of each image because the time label and scale bar are automatically superimposed on the IncuCyte ZOOM™ images and these objects partially obscure the numbers of cells in these subregions of each images. We also discard the left-most column from each image, which leaves us with a slightly smaller image that is discretized into 37 equally-spaced columns of width 49.5 *μ*m, and a reduced domain width of 1831 μm. Next, to reduce the magnitude of the fluctuations in the cell density profile, we average the density profiles associated with each of the four experimental replicates to obtain a single averaged cell density profile as a function of time, as summarized in Fig 6. For completeness, the relatively noisy column-averaged cell density profiles associated with each of the four individual experimental replicates are given in the Supporting Information (S2 Fig). Visual inspection of the averaged density profiles in Fig 6 confirms that we observe a well-defined vacant region at the beginning of the experiment and that we see the vacant region become smaller with time as the two opposingly-directed cell fronts move to close the vacant region over time.

**Figure 6:**
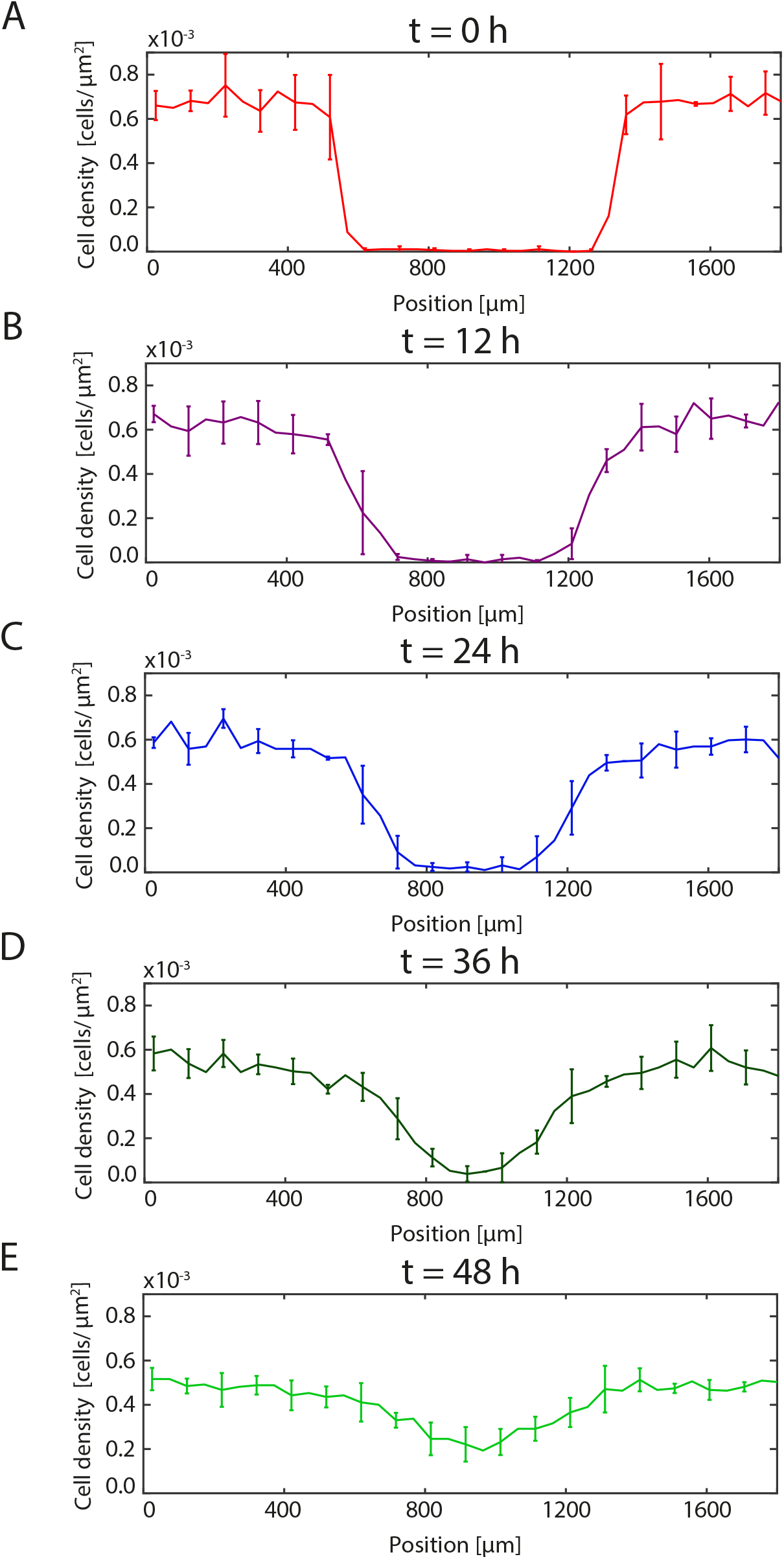
Averaged cell density profiles. A-E: Averaged cell density profiles at *t* = 0, 12, 24, 36, and 48 h, as indicated. All profiles report the sample mean density computed using four identically prepared experimental replicates. The error bars denote the sample standard deviation. For clarity we only present the error bar at every second column.

Now that we have quantified our experimental observations in terms of the temporal variation in the column-averaged cell density profiles, further averaged across four identically prepared experimental replicates, we will now attempt to use a suite of discrete mathematical models to mimic the experimental data set. To provide the most realistic discrete simulations, we always take care to initialize each discrete simulation using the exact same number and locations of cells that are present in the experimental images at *t* = 0 h. To do this we extract the position of each individual cell from the experimental images at *t* = 0 h using the object identification tool in CellProfiler, Fig 5. This approach means that our initial condition for the discrete simulations is precisely the same as in our experimental data set.

### Discrete stochastic model

To simulate our experimental data set we use a novel two-dimensional discrete model of cell motility that incorporates random motility, intercellular interactions including both long range cell-to-cell adhesion and short range cell pushing, as well as capturing dynamic changes in cell size. We choose to work with a discrete modelling framework because discrete individual-based approaches are more natural to compare with experimental images than continuum models. Such discrete individual-based models are used to study a range of cell biology phenomena including, malignant invasion [40], wound healing [41], self-organization [42], angiogenesis [43] and embryonic development [13]. Since we work with an off-lattice discrete framework, each agent is allowed to move in any direction on a continuous domain. This off-lattice approach is more realistic than a simpler lattice-based model where the locations of agents are restricted to an artificial lattice structure [18, 44, 45, 42, 46].

Our modelling approach is flexible since it can incorporate a range of individual-level processes such as random motility, long range cell-to-cell adhesion, short range cell pushing and captures dynamic changes in cell size. We refer to this model as *Model I* since it describes the most fundamental situation where all four processes are acting simultaneously. We begin by introducing Model I and then describe three simplifications of this model in which we systematically neglect certain features. These simplifications include: (i) capturing random motility, long range cell-to-cell adhesion and short range cell pushing (*Model II*); (ii) capturing long range cell-to-cell adhesion, short range pushing *(Model III*); and (iii) capturing random motility only (*Model IV*). To explore the relevance of the mechanisms inherent in these four models we carefully calibrate each model to match the density data summarized in Fig 6 and quantitatively compare the results of the calibration procedure.

### Model I: random motility, long range cell-to-cell adhesion, short range cell pushing and dynamic changes in cell size

A key novelty of our approach is that we simulate the dynamic change in agent diameter, *δ*(*t*). This approach is very different to standard approaches where agents in discrete models are thought of as having either no size or a constant size. Here, to match our experimental measurements presented in Fig 2(C), we assume that *δ*(*t*) increases linearly,

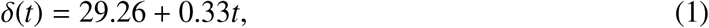

where *t* is time, measured in hours.

We assume that the location of each individual in the population is described by an appropriate equation of motion [42, 47, 48], and we employ a Langevin stochastic framework where a population of *N* cells is modelled by a system of *N* first order stochastic differential equations

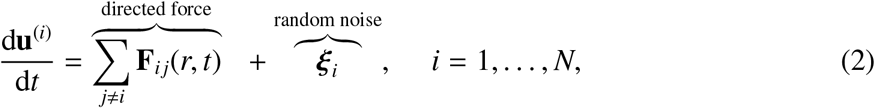

where **u**^(*i*)^ is the position vector of the ith agent on a two-dimensional domain, **F***_ij_* is the deterministic interaction force between agents *i* and *j* that are separated by distance *t*, and ***ξ**_i_* is a random stochastic force exerted on the ith agent. The stochastic ***ξ**_i_* force is sampled from the Gaussian distribution [48] with zero mean and a variance, 2*D*/Δ*t*, where *D* is the diffusivity, and Δ*t* is the time step used to numerically solve Eq 2. Since the Langevin equation formalism does not include any inertial forces, this framework implicitly neglects agent acceleration. This assumption is reasonable at low Reynolds numbers, and is routinely invoked at cellular length scales [49].

The Langevin stochastic model, Eq 2, implies that the movement of each agent is determined by the combined effects of the deterministic intercellular force, **F***_ij_*, and the random stochastic force, ***ξ**_i_*. The details of the deterministic interaction force, **F***_ij_*, can be chosen to incorporate a range of relevant phenomena such as long range attraction and short range repulsion [50], as illustrated in Fig 7. In this work we specify that the interaction force, **F***_ij_*, depends upon the distance between agents, *r*, and time, *t*, and is given by

**Figure 7:**
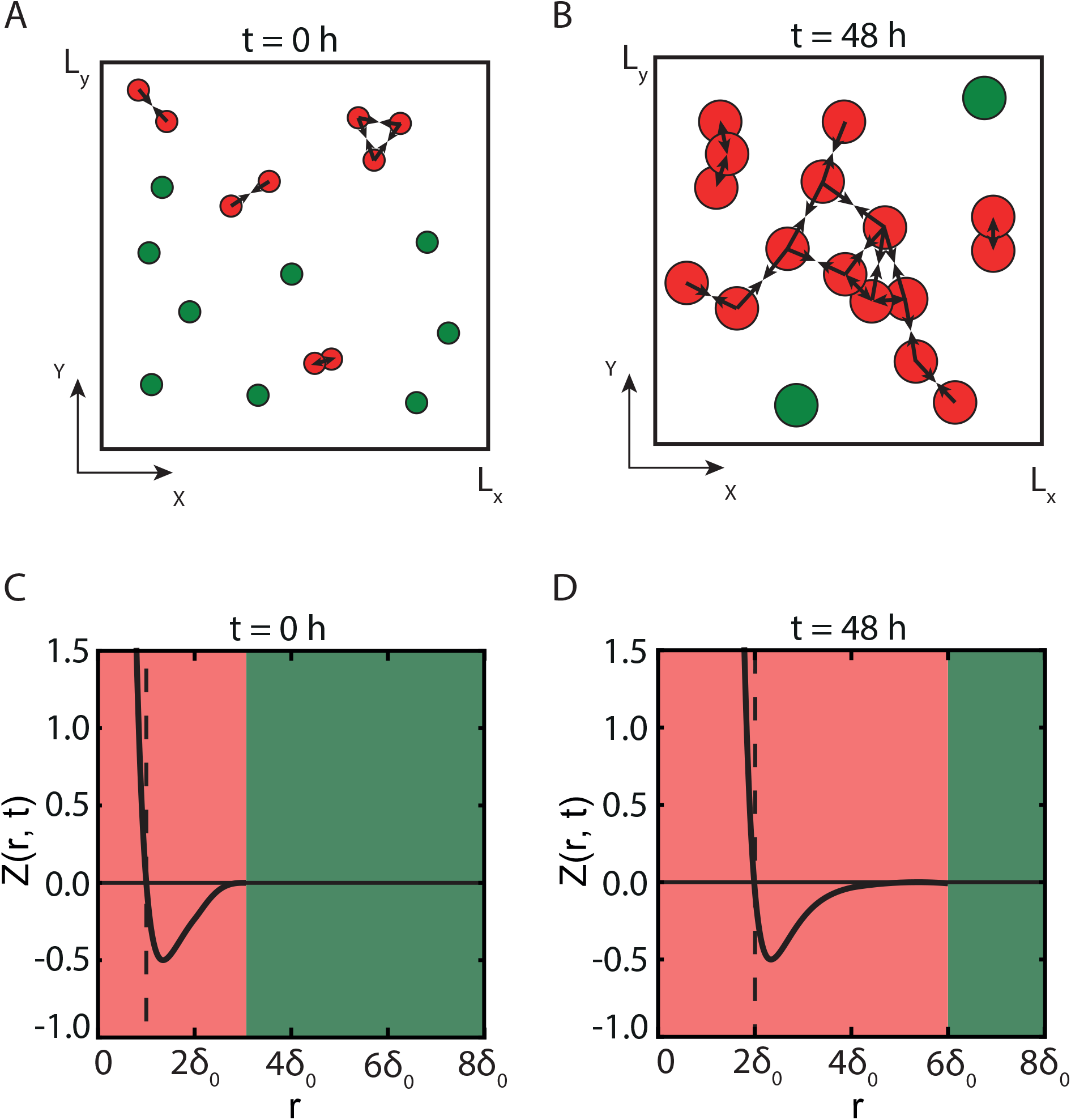
Schematic showing how short and long range forces are introduced in the model through the dimensionless force function, *Z*(*r*, *t*). A-B: Green circles denote isolated agents that are unaffected by cell-to-cell interactions and, as a consequence, undergo random migration. Red circles indicate agents that are sufficiently close to other agents that they interact with them. Comparing the schematics in A and B shows that the increase in agent size leads to additional agent-to-agent interactions because agent growth reduces the distance between agents. The arrows in A and B indicate the direction of the deterministic forces **F***_ij_*. C-D: Dimensionless force function *Z*(*r*, *t*) for *t* = 0 h and *t* = 48 h. *δ*_0_ is the agent size at the start of the experiment, *t* = 0 h. The red shaded regions indicate a sufficiently small distance between agents, *r* ≤ 3*δ*(*t*), where agent-to-agent interactions are present. The green shaded regions indicate a sufficiently large distance between agents, *r* > 3*δ*(*t*), so that there is no interaction.

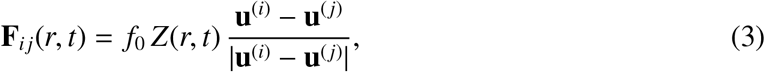

where *f*_0_ is dimensional magnitude of the interaction force, *Z*(*r*, *t*) is a dimensionless function describing how the interaction force depends on the separation between the agents, *r* = |**u**^(*i*)^ − **u**^(*j*)^|. As with all models of the form of Eq 2, the dimensional magnitude of the interaction force, *f*_0_, has dimensions of velocity.

In this work we use the dimensionless function, *Z*(*r*, *t*), to incorporate three main features of cell-cell interactions: (i) short range repulsion forces which mimic cell pushing; (ii) longer range attraction forces which mimic cell-to-cell adhesion; and (iii) dynamic changes in agent size. The short range repulsion forces can be interpreted as cell resistance to deformation, which leads to crowding and volume exclusion effects [51]. In contrast, the longer range attraction forces mimic intercellular attraction. These cell-to-cell attraction forces are thought to be a predominant factor in the cell-cell adhesion [52]. To incorporate these effects we use a modified Morse potential [53],

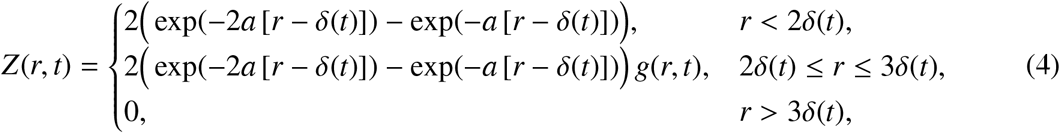

where *δ*(*t*) is the time-dependent agent diameter, *a* > 0 is a parameter that controls the shape of the force function, and *g*(*r*) is the Tersoff cut-off function [54] which ensures a finite range of interactions,

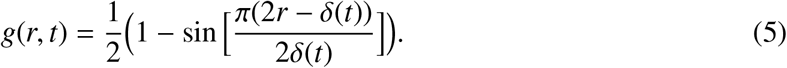

The cell-cell interaction range is finite, and set to three agent diameters [47]. As such, the interaction force is zero for separation distances of greater than three agent diameters. For all results presented here we set *a* = 0.08 which implies that the agents are relatively rigid and highly unlikely to overlap [53].

Schematics showing the key features of *Z*(*r*, *t*) at *t* = 0 h and *t* = 48 h are shown in Fig 7(C-D). Here we see that at short separation distances we have strong positive *Z*(*r*, *t*), which captures short range repulsion and pushing, owing to finite size effects. Over longer separation distances we have smaller negative *Z*(*r*, *t*) which models attraction, such as adhesion. Finally, over sufficiently large distances we have no interactions as *Z*(*r*, *t*) = 0. To capture the effects of the increase in cell size, all length scales in Fig 7(C-D) are given in terms of the average cell diameter, *δ*(*t*), which can vary with time. In this work we use a linear function for *δ*(*t*) because this matches our experimental observations, however other functional forms for *δ*(*t*) are possible, for example if we were to consider experiments performed over a longer time scale where the linear increase would not be relevant.

In all simulations we apply the Langevin model on a domain of size 1831.5 *μ*m × 1439 *μ*m, which is the size of the experimental field of view after the boundary columns have neglected. Furthermore, we always initiate each stochastic simulation so that the initial number and location of agents in the simulation corresponds to the initial number and location of cells in the experiments, as shown in Fig 5. Fig 8 shows a schematic of how we apply this model to simulate the geometry of the IncuCyte ZOOM™ assay. In this schematic we highlight the spatial arrangement of agents in the simulation and the interaction forces acting upon these agents.

**Figure 8:**
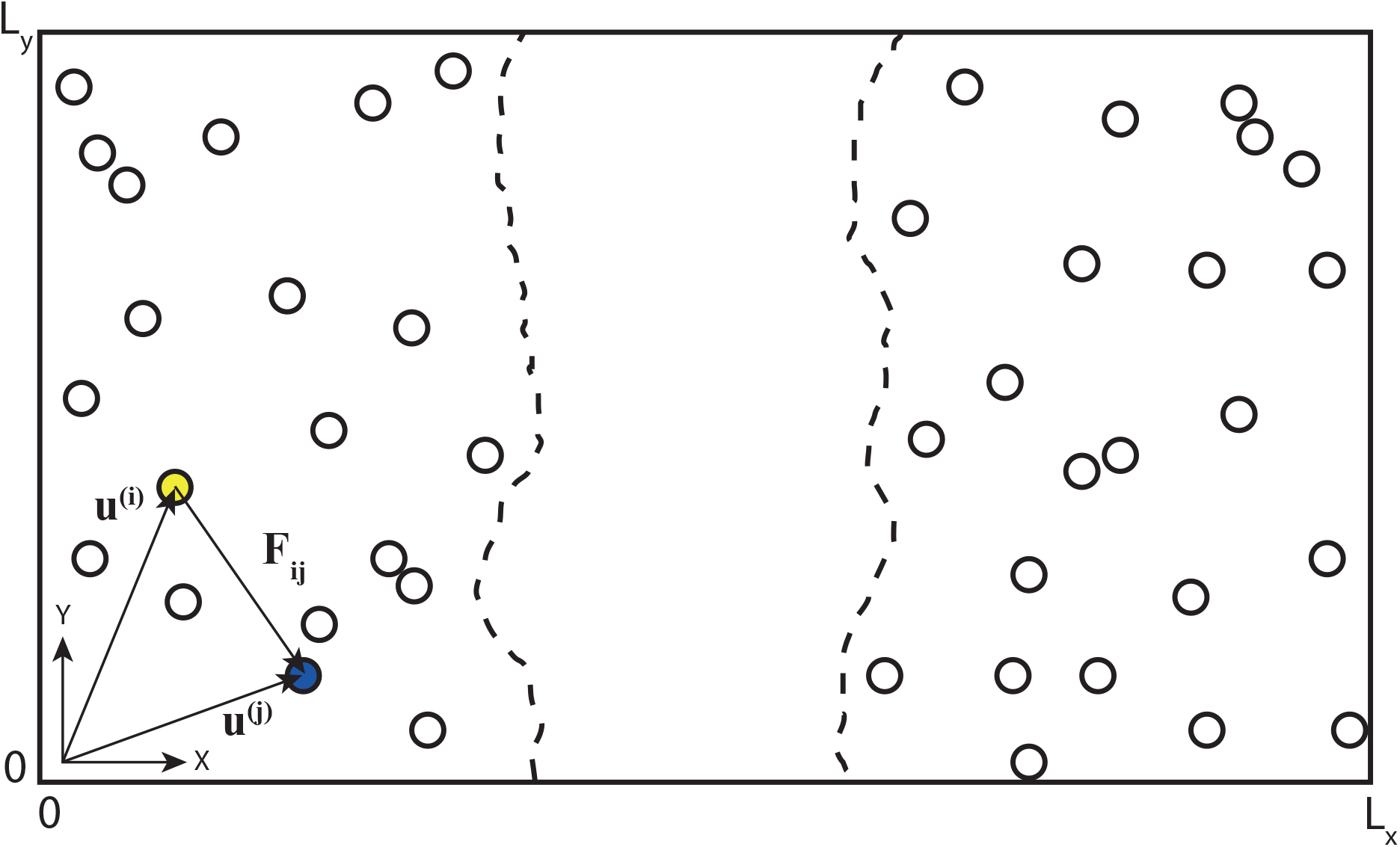
Schematic of the experiment. Vectors **u**^(*i*)^ and **u**^(*j*)^ denote position vectors of two arbitrary agents. Vector **F***_ij_* is the interaction force between the yellow agent *i* and the blue agent *j*. Dashed lines represent the approximate boundary between the initially-vacant and the initially-occupied regions.

### Model II: random motility, long range cell-to-cell adhesion, short range cell pushing and constant cell size

Model II differs from Model I only in that we make the standard assumption that the size of the cells remain constant during the cell migration process. Consequently, we set *δ*(*t*) = *δ*_0_ = 29.26 *μ*m during the simulations. This number corresponds to the average diameter of the cells at the beginning of experiments, *t* = 0 h. All other features of Model II are identical to Model I.

### Model III: long range cell-to-cell adhesion, short range pushing, dynamic cell size changes and no random motility

Model III differs from Model I only in that we neglect the role of random motility. Therefore, setting ***ξ**_i_* = 0 we rewrite Eq 2 as

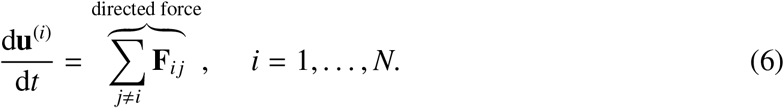

It is useful to note that with a deterministic initial condition, the solution of Eq 6 is deterministic.

### Model IV: random motility only

Model IV differs from Model 1 in that we neglect the role of the interaction force and cell migration is purely random, implying that agents are point particles without any size or deterministic interactions. This assumption reduces the system of Langevin equations to a Brownian process which can be written as

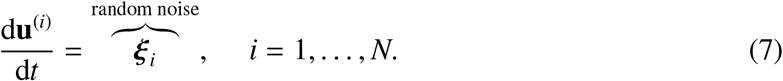

As we described previously, the initial coordinates of agents in each stochastic simulation is taken to be the precise position of each individual cell at *t* = 0 h, as depicted in Fig 5. To simulate Models I–IV we must apply appropriate boundary conditions to reflect the conditions relevant to the experiment. To determine these boundary conditions, we note that the experimental images, shown in Fig 2(A-B), correspond to a field of view that is much smaller than the spatial extent of the experiment. For example, the width of the field of view in Fig 2(A-B) is 1979 *μ*m, which is much smaller than the diameter of the well in the 96-well plate, which is 9000 *μ*m, as shown in Fig 3. The schematic in Fig 3 is important because it emphasizes that the images from this kind of assay only show a small proportion of the population of cells present in the well. In particular, it is important to remember that the boundaries around the field of view are not physical boundaries since the spatially uniform population of cells extends far beyond the boundaries around the field of view. This means that whenever the density is below confluence, cells will migrate, in each direction, across the boundaries of the field of view. However, since the population of cells is placed uniformly into each well of the 96-well plate, the net flux of cells across the boundaries of the field of view will be zero owing to symmetry. Therefore, the appropriate boundary conditions for all of our simulations is to impose zero net flux boundary conditions around all boundaries of the field of view [55]. We implement these boundary conditions by simply aborting any potential movement event that would take a particular agent across one of the boundaries.

All model simulations presented in this work are obtained by numerically integrating the governing equations using a forward Euler method with a constant time step of duration *δt* = 0.02 h. This choice of time step is sufficiently small that our results are grid-independent.

## Results and Discussion

To quantitatively compare the suitability of Models I–IV to describe our experimental data set, we calibrate each model to provide the best match to the experimental density data. For Model 1 our aim is to estimate the two model parameters, (*D*, *f*_0_), that leads to the best match with the experimental measurements. To facilitate this we introduce a measure of the discrepancy between the experimental data and the model solution,

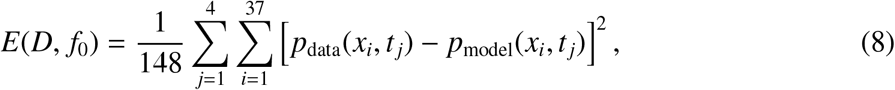

where *E*(*D*, *f*_0_) measures the discrepancy between the experimental cell density, *p*_data_(*x_i_*, *t_j_*), and the density predicted by the model, *p*_model_(*x_i_*, *t_j_*), for given values of *D* and *f*_0_. The index *i* denotes column number along *x* coordinate, and index *j* denotes time so that *j* = 1, 2, 3 and 4 correspond to the experimental time points *t* = 12, 24, 36, and 48 h, respectively. The experimental density estimates, *p*_data_(*x_i_*, *t*), corresponds to the averaged experimental density, where the average is taken across all four identically-prepared experimental replicates. We find that it is necessary to estimate *E*(*D*, *f*_0_) using averaged experimental data, rather than working with the four experimental data sets separately since the fluctuations in the data from the individual replicates lead to sufficiently large fluctuations in our estimates of *E*(*D*, *f*_0_). Throughout this work we will refer to the function *E*(*D*, *f*_0_) as an *error surface* as we will visualize the surface and seek to find values of *D* and *f*_0_ that minimize the error, or discrepancy between the experimental density data and the density data predicted by the mathematical models. Overall, our aim is to find estimates of *D* and *f*_0_ that minimize *E*(*D*, *f*_0_), and we denote these estimates as 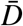 and 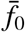, respectively. All results, for Models I–IV, are obtained by simulating the stochastic models in two-dimensional space. We then construct one-dimensional density distributions from the model output, *p*_model_(*x_i_*, *t_j_*), using the same procedure that is used to convert the distribution of cells in the two-dimensional experimental images into one-dimensional density profiles.

Since we have access to four identically prepared initial conditions for our experimental data set, each time we attempt to match the models with the averaged experimental data we repeat the process four times using the four different choices of initial conditions associated with experimental data sets A, B, C and D. This approach means that we can estimate and visualize the error surface four times for each particular model. Despite the additional computational overhead associated with this approach, we note that it is a useful approach because it provides us with additional quantitative insight about the suitability of each model. In particular, a mathematical model that is compatible with the data ought to lead to estimates of *D* and *f*_0_ that are consistent across the four initial conditions, and so we will examine Models I–IV to see whether they are capable of providing parameter estimates that are consistent across the four different initial conditions.

Previous modelling studies based on using reaction-diffusion equations indicate that estimates of diffusivity for PC-3 prostate cancer cells varies significantly, from about 300 *μ*m^2^/h for low density conditions to approximately 1000 *μ*m^2^/h for high density conditions [56]. Consequently, we focus our search for parameter estimates in the interval 0 ≤ *D* ≤ 1100 *μ*m^2^/h. In contrast, there are no previous estimates of *f*_0_ for the PC-3 cell line. Since we have little initial guidance about an appropriate choice of *f*_0_, we first conducted a series of preliminary simulations (not shown) to determine an acceptable range of *f*_0_ for each model. This exercise suggests that acceptable ranges are approximately 0 ≤ *f*_0_ ≤ 0.11 *μ*m/h for Model I; 0 ≤ *f*_0_ ≤ 0.6 *μ*m/h for Model II; and 0 ≤ *f*_0_ ≤ 0.11 *μ*m/h for Model III.

To estimate 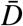 and 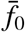 for Model I we estimate discrete values of *E*(*D*, *f*_0_) by using a series of numerical solutions of Eq 2 over many (*D*, *f*_0_) pairs. For each parameter pair we generate an ensemble of 100 identically prepared realizations of the stochastic model and then estimate *E*(*D*, *f*_0_) by averaging the density data from the 100 identically prepared realizations. Results in Fig 9(A-D) show contour plots of the error surface constructed using seven values of diffusivity, *D* = 0, 100, 300, 500, 700, 900, 1100 *μ*m^2^/h, and 12 equally-spaced values of the force amplitude in the interval 0 ≤ *f*_0_ ≤ 0.11 *μ*m/h. All contour lines in Fig 9 are obtained using Matlab’s spline interpolation function griddedlnterpolant. The four different contour plots in Fig 9(A-D) show separate error surfaces obtained by applying Model I to each of the four identically prepared experimental replicates. Since the initial positions of agents in the discrete simulations exactly correspond to the positions of cells in experimental images, given in Fig 5, the magnitude of the fluctuations in the experimental data are consistent with the magnitude of the fluctuations from one realization of the stochastic model. In general, the magnitude of the fluctuations in the stochastic model scales with the number of realizations of the stochastic model, 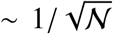. Therefore, our choice of using 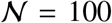 realizations leads to averaged discrete density profiles with fluctuations that are approximately one order of magnitude smaller than fluctuations in the experimental data.

**Figure 9:**
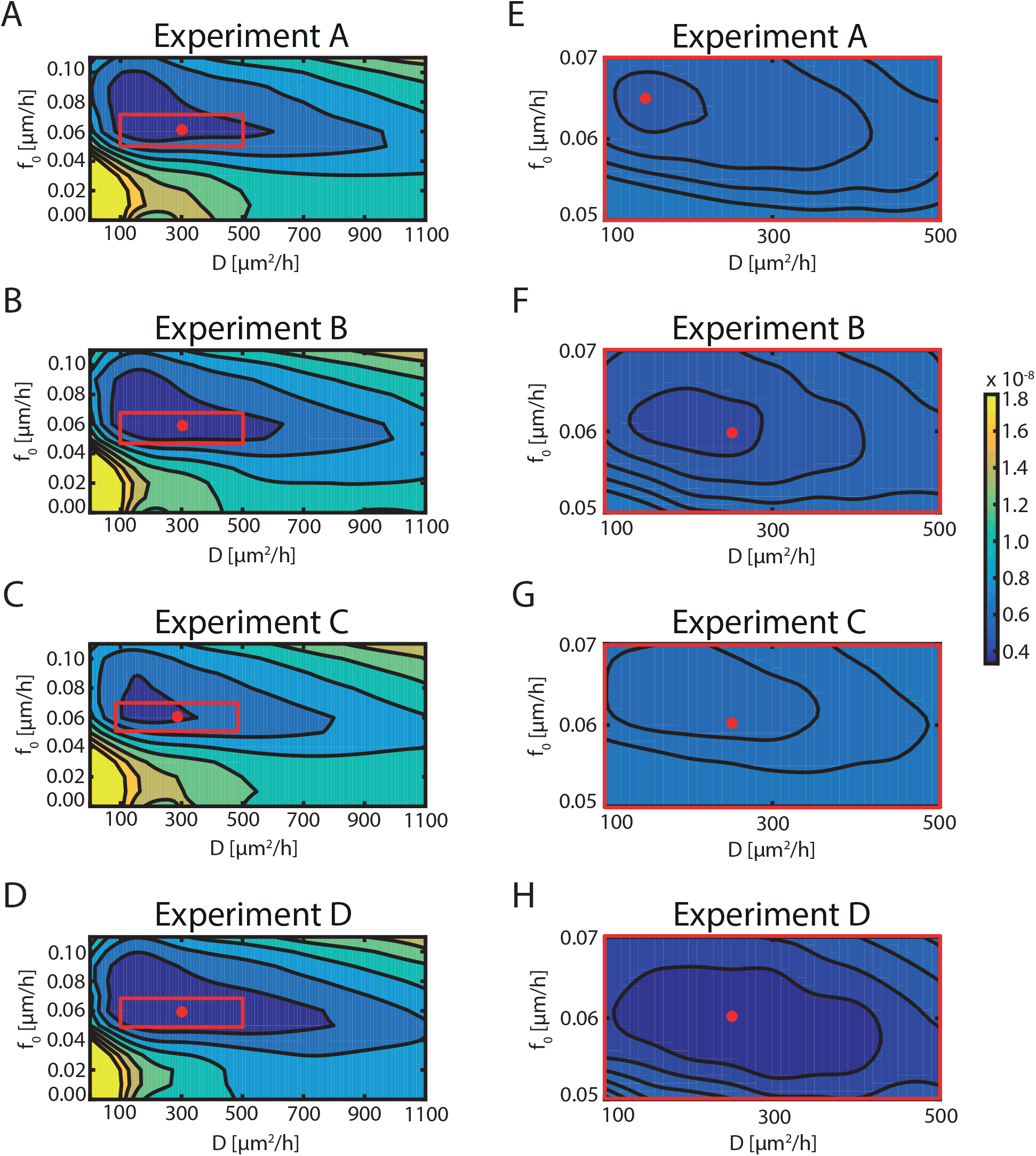
*E*(*D*, *f*_0_) for Model I. A-D: Error surface contours, *E*(*D*, *f*_0_), for four identically-prepared IncuCyte ZOOM™ scratch assays. Each surface contour plot is constructed using seven values of the diffusivity in 0 ≤ *D* ≤ 1100 *μ*m^2^/h, and 12 equally-spaced values of the force amplitude in 0 ≤ *f*_0_ ≤ 0.11 *μ*m/h. E-H: Refined error surface contours obtained by estimating *E*(*D*, *f*_0_) on a refined discretisation of the parameter space within the red rectangles in A-D. The values of *E*(*D*, *f*_0_) are shown on the color bar. The location of the best-fit estimate 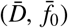 is shown as a red circle in each subfigure.

Preliminary estimates of 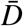 and 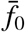 are obtained by evaluating the error surface across a relatively coarse discretisation of the parameter space. These estimates are shown in Fig 9(A-D) as red circles. We then refine our estimates of 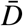 and 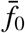 by considering a refined discretisation of a subregion surrounding each red circle in Fig 9(A-D). These subregions are shown as red rectangles in Fig 9(A-D). Refined plots of the error surface in Fig 9(E-H) are obtained by calculating *E*(*D*, *f*_0_) across five equally-spaced values of *D* in 100 ≤ *D* ≤ 500 *μ*m^2^/h and five equally-spaced values of *f*_0_ in 0.05 ≤ *f*_0_ ≤ 0.07 *μ*m/h. Each individual plot of *E*(*D*, *f*_0_) in Fig 9(E-H) shows that we have a well-defined minmum from which we can estimate 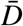 and 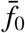. Comparing data in Fig 9(E-H) shows that our estimates of 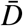 and 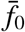 are consistent between the four different experimental replicates. In fact, three of the four refined plots give remarkably consistent estimates of 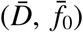. Only one of the experimental replicates, shown in Fig 9(E), gives slightly different parameter estimates, 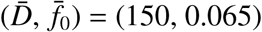.

To visualize the quality of match between the experimental data and discrete profiles predicted by the stochastic model we use the best-fit parameter estimates 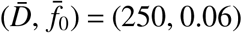 for experiment D. First, we solve Model I with 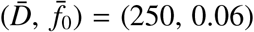 and calculate the density profiles from the simulations as before. Second, we superimpose density profiles from the discrete simulations with the experimental density distributions, as shown in Fig 10. The choice of initial conditions in the stochastic model guarantees that we have an exact match between the experimental density profile and the simulation density profiles at *t* = 0 h. However, since we are dealing with stochastic experimental data and a stochastic mathematical model we do not expect there to be an exact match at later times. Results in Fig 10(A-B) show that we have a reasonable match between the calibrated simulation results and the experimental density data for a single realization of the stochastic model.

**Figure 10:**
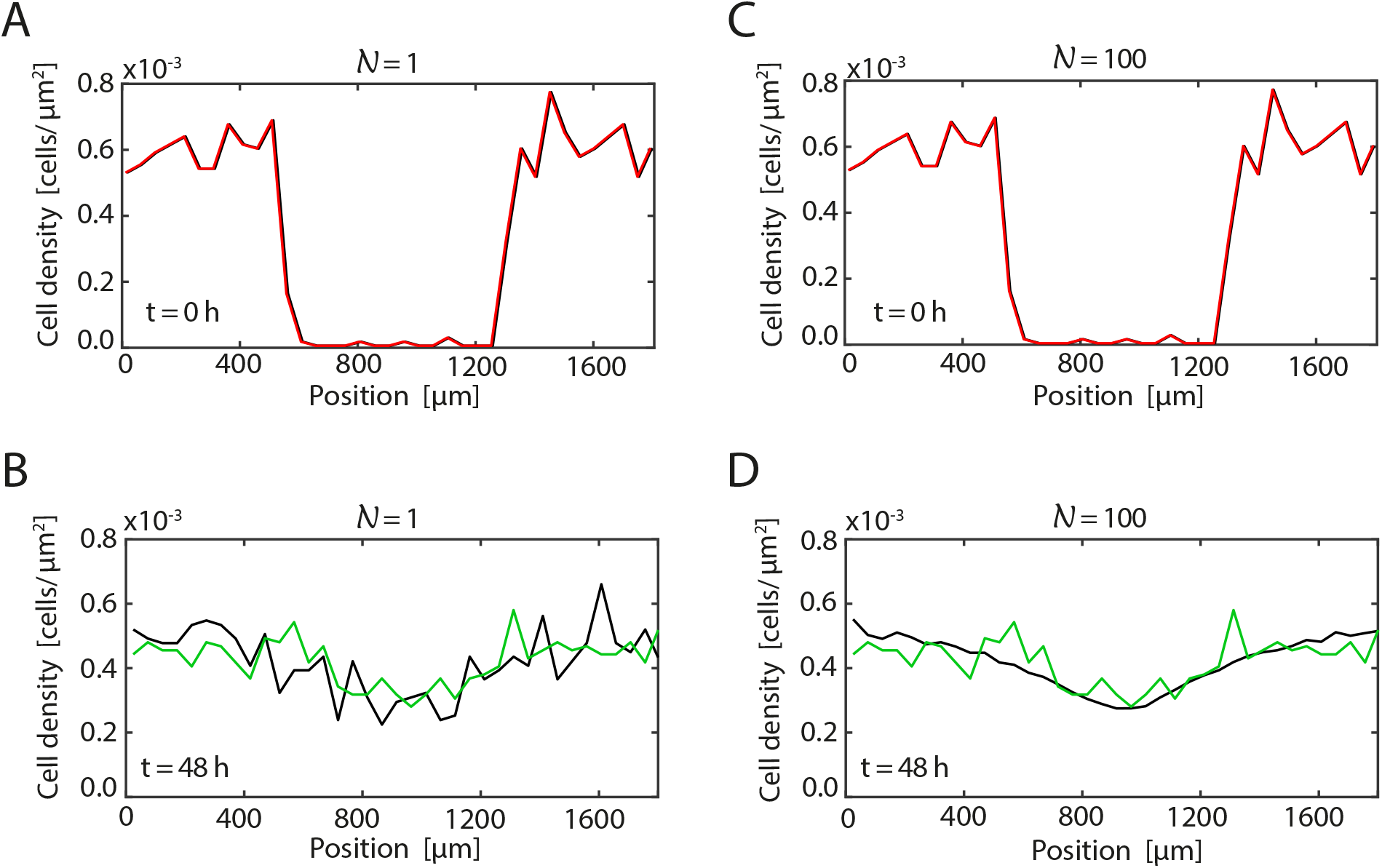
Comparison of the calibrated solutions of Model I and experimental profiles from experiment D. Results in A-B compare the experimental data (red, green) and Model I solution (black) at *t* = 0 and *t* = 48 h, respectively, for 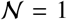 model realization. Results in C-D compare the experimental data (red, green) and Model I solution (black) at *t* = 0 and *t* = 48 h, respectively, averaged over 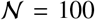 identically prepared realizations of the model. The best-fit parameter estimates are 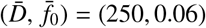 from Fig 9(H).

Similarly, results in Fig 10(C-D) show that we also have a good match between the simulation results and the experimental density data over 100 identically prepared realizations of the stochastic model where we see that the magnitude of the fluctuations in the averaged stochastic data are reduced.

We now turn our attention to calibrating Model II to match the experimental data. To achieve this we repeat the exact same calibration process except that we implement Model II with constant agent size. We repeat the process of generating error surfaces using Model II. Comparing results in Fig 9(A-D) and Fig 11(A-D) shows that the best fit parameters in Model II lead to a larger value of *E*(*D*, *f*_0_). Furthermore, comparing estimates of 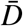 and 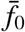 for Model II between the four identically prepared experimental data sets shows that we have a much higher degree of variability between our parameter estimates for Model II than we did for Model I. Overall, these results suggest that Model I is more consistent with our experimental data than Model II, and so we do not proceed with any further refinement of our parameter estimates for Model II. This result shows that the traditional approach of neglecting the dynamical changes in cell size has clear impact on the ability of the model to describe the behaviour of the entire cell population.

To visualize the quality of match between the experimental data and best-fit density profiles predicted by Model II we use the best-fit parameter estimates 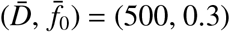 for experimental replicate D. Again, we solve Model II with these parameter estimates and then estimate the density profiles from those simulations. Results in Fig 12 show the density profiles from the discrete simulations superimposed on the corresponding experimental density distributions. Visually, we see that the quality of match in Fig 12(B) and Fig 12(D) is notably poorer than the quality of match in Fig 10(B) and Fig 10(D). This visual difference is consistent with the quantitative differences in *E*(*D*, *f*_0_) in Fig 9 and Fig 11.

**Figure 11:**
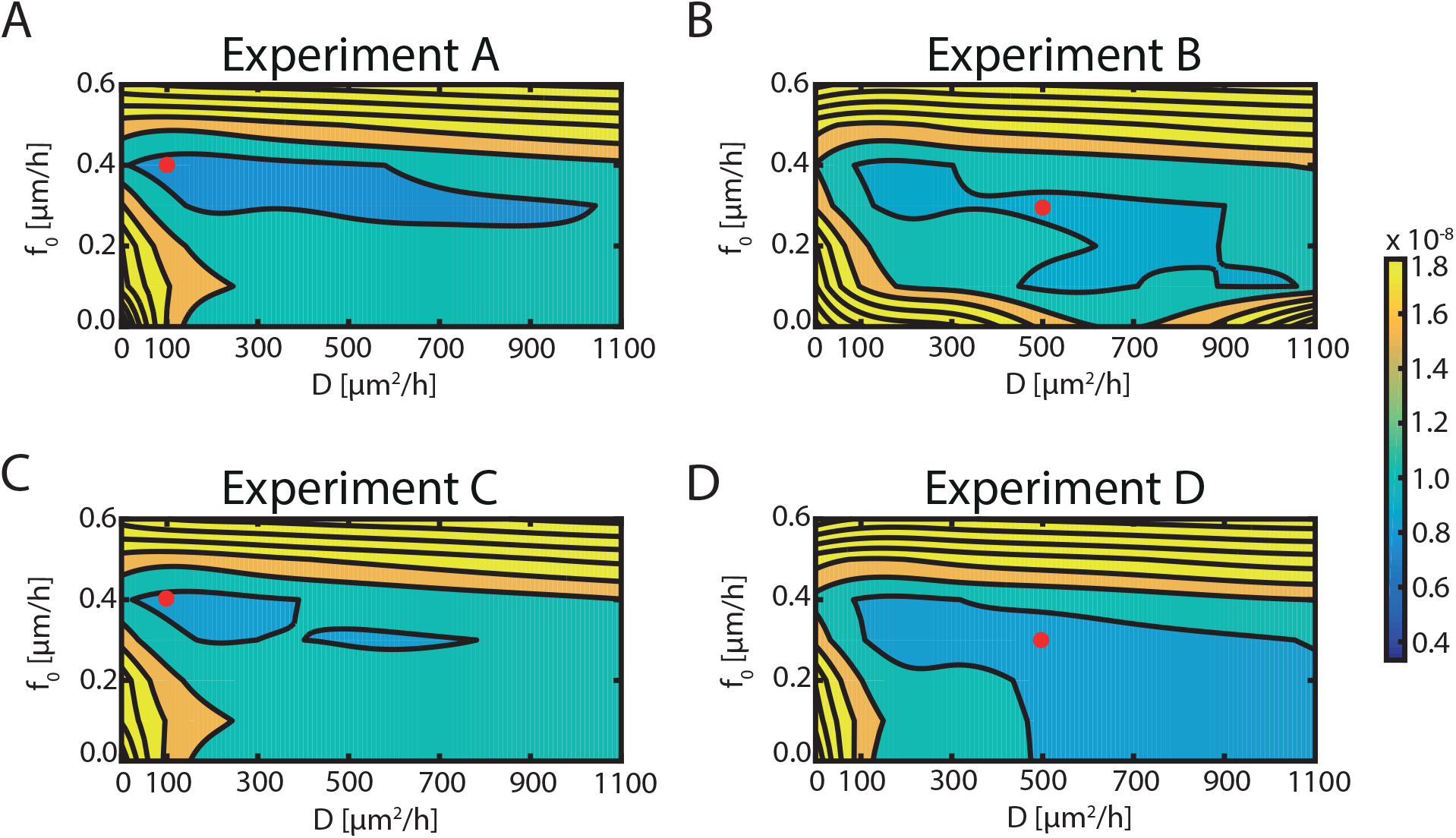
*E*(*D*, *f*_0_) for Model II. A-D: Error surface contours, *E*(*D*, *f*_0_), for four identically-prepared IncuCyte ZOOM™ scratch assays. Each surface contour plot is constructed using seven values of diffusivity in 0 ≤ *D* ≤ 1100*μ*m^2^/h and seven equally-spaced values of the force amplitude in 0 ≤ *f*_0_ ≤ 0.6 *μ*m/h. The values of *E*(*D*, *f*_0_) are shown on the color bar. In each case the location of the best-fit estimate 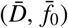 is shown as red circle.

**Figure 12:**
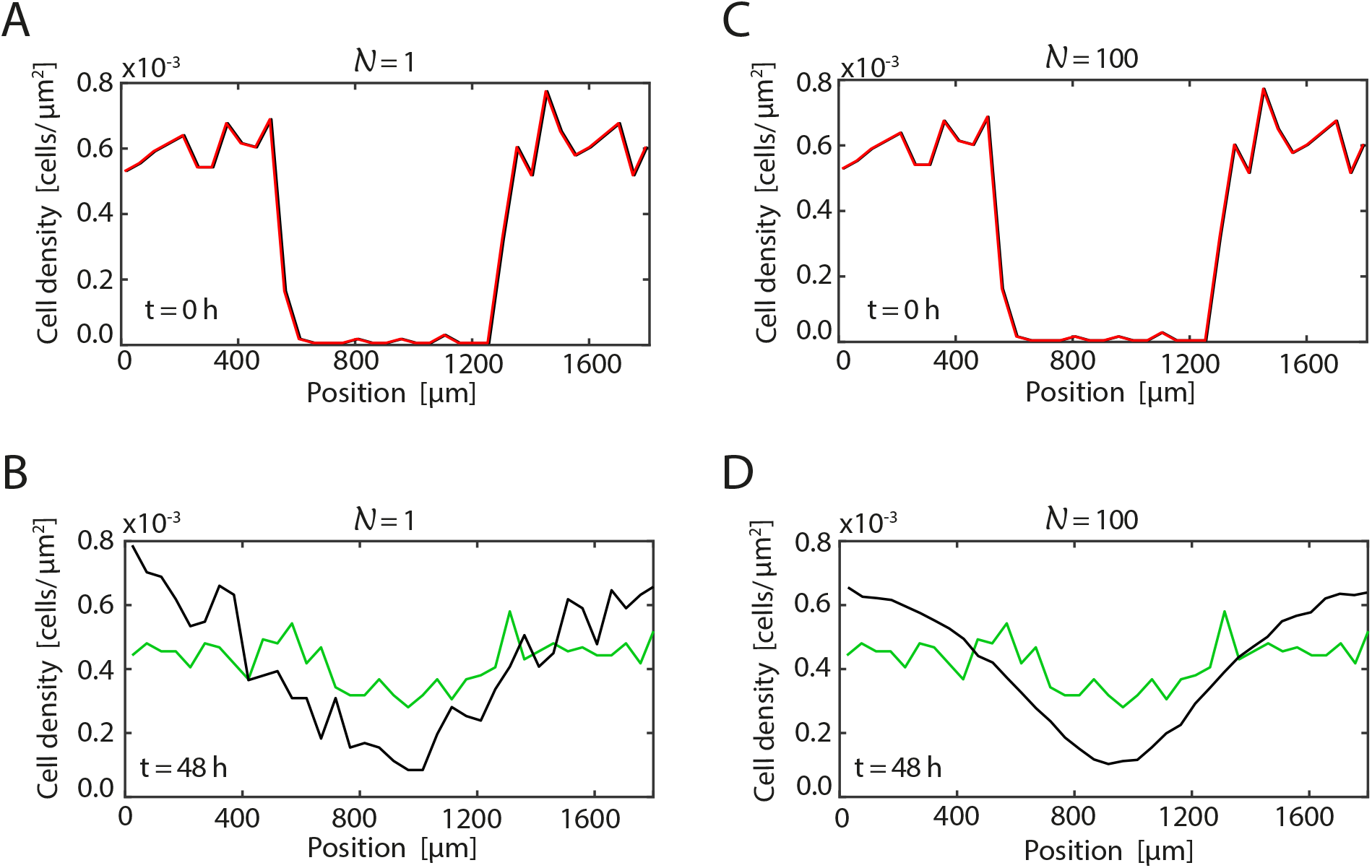
Comparison of the calibrated solutions of Model II and experimental profiles from experiment D. Results in A-B compare the experimental data (red, green) and Model II solution (black) at *t* = 0 and *t* = 48 h, respectively, for 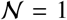 model realization. Results in C-D compare the experimental data (red, green) and Model II solution (black) at *t* = 0 and *t* = 48 h, respectively, averaged over 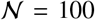 identically prepared realizations of the model. The best-fit parameter estimates are 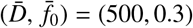 from Fig 11(D).

Finally, we calibrate Models III and IV to match the experimental data. Note that Model III neglects the role of random motility so our parameter estimation involves just one parameter, *f*_0_. Similarly, Model IV neglects the role of intercellular forces so our parameter estimation involves just one parameter, *D*. Following a now familiar procedure, we compute measures of discrepancy, *E*(*f*_0_) and *E*(*D*), for Models III and IV, respectively. Results in Fig 13(A) show a well-defined minimum for each of the four experimental replicates for Model III, however the best-fit estimates are in the range 0.07 ≤ *f*_0_ ≤ 0.1 *μ*m/h, which is approximately double the estimate we identified previously for Model I. In contrast, results in Fig 13(B) show that we have relatively poorly-defined minimum for all four experimental replicates for Model IV. In this case the best-fit estimates are in the range 700 ≤ *D* ≤ 1000 *μ*m^2^/h which is approximately four times greater than the estimates we obtained for Model I.

**Figure 13:**
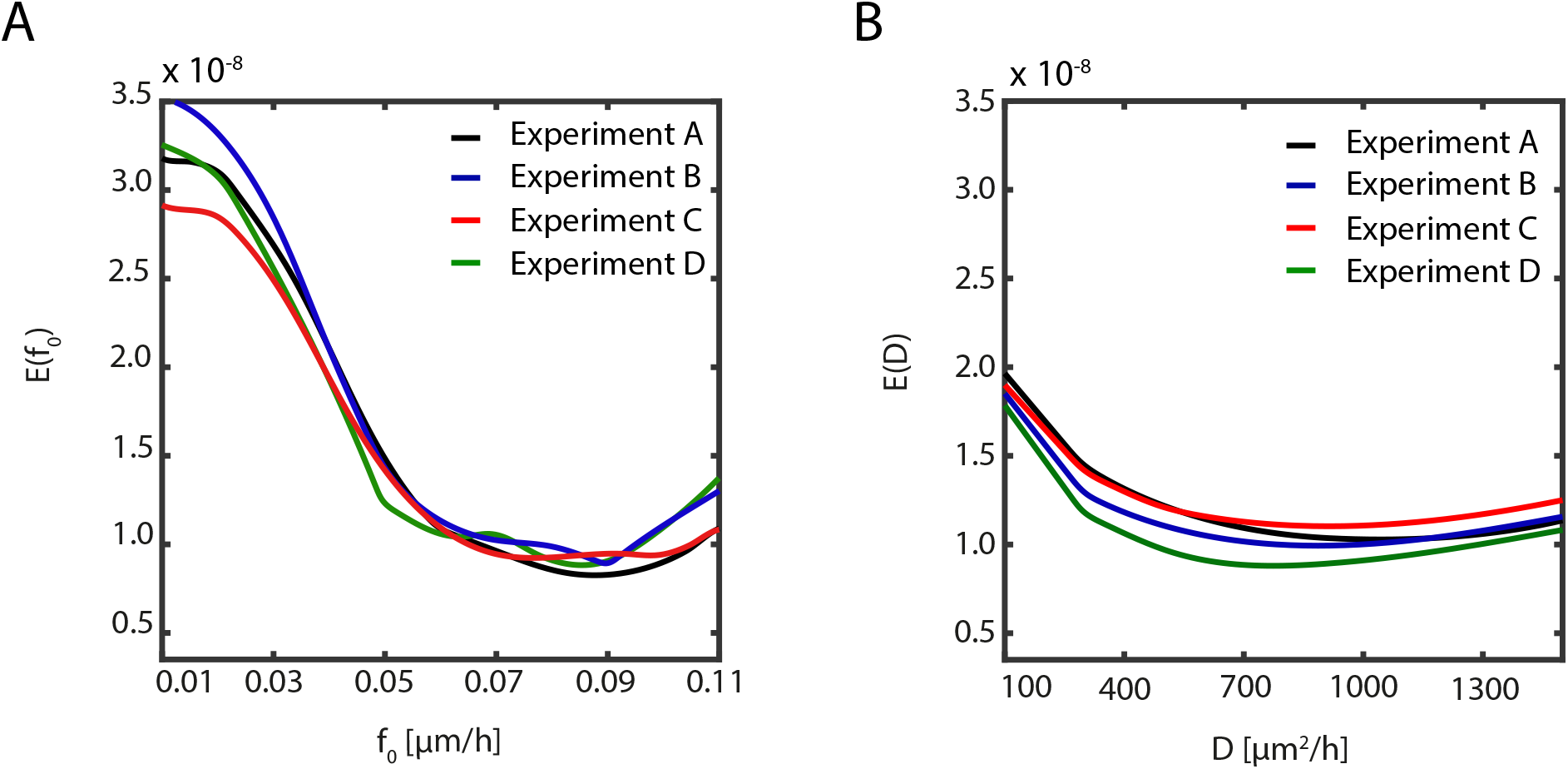
*E*(*f*_0_) and *E*(*D*) for Models III and IV. A: *E*(*f*_0_) for Model III constructed for each experimental replicate with 11 equally-spaced values of *f*_0_ in 0.01 ≤ *f*_0_ ≤ 0.11 *μ*m/h. B: *E*(*D*) for Model IV constructed for each experimental replicate with eight equally-spaced values of *D* in 100 ≤ *D* ≤ 1500 *μ*m^2^/h.

Results in Fig 14 compare the discrete density profiles obtained using Model III parameterized with the best-fit estimate *f*_0_ = 0.09 *μ*m/h and the experimental density profiles from experiment D. We note that Model III is deterministic and produces the same density distribution regardless of the number of model realizations. The quality of match between calibrated Model III and the experimental data for experimental replicate D is reasonable, however the value of 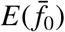 is greater than the value of 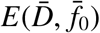 for Model I, thereby indicating that Model I produces an improved match to the experimental data. Results in Fig 15 compares discrete density profiles obtained using Model IV parameterized with the best-fit estimate *D* = 700 *μ*m^2^/h where we see no improvement in the quality of match between the calibrated mathematical model and the experimental data relative to Model 1.

**Figure 14:**
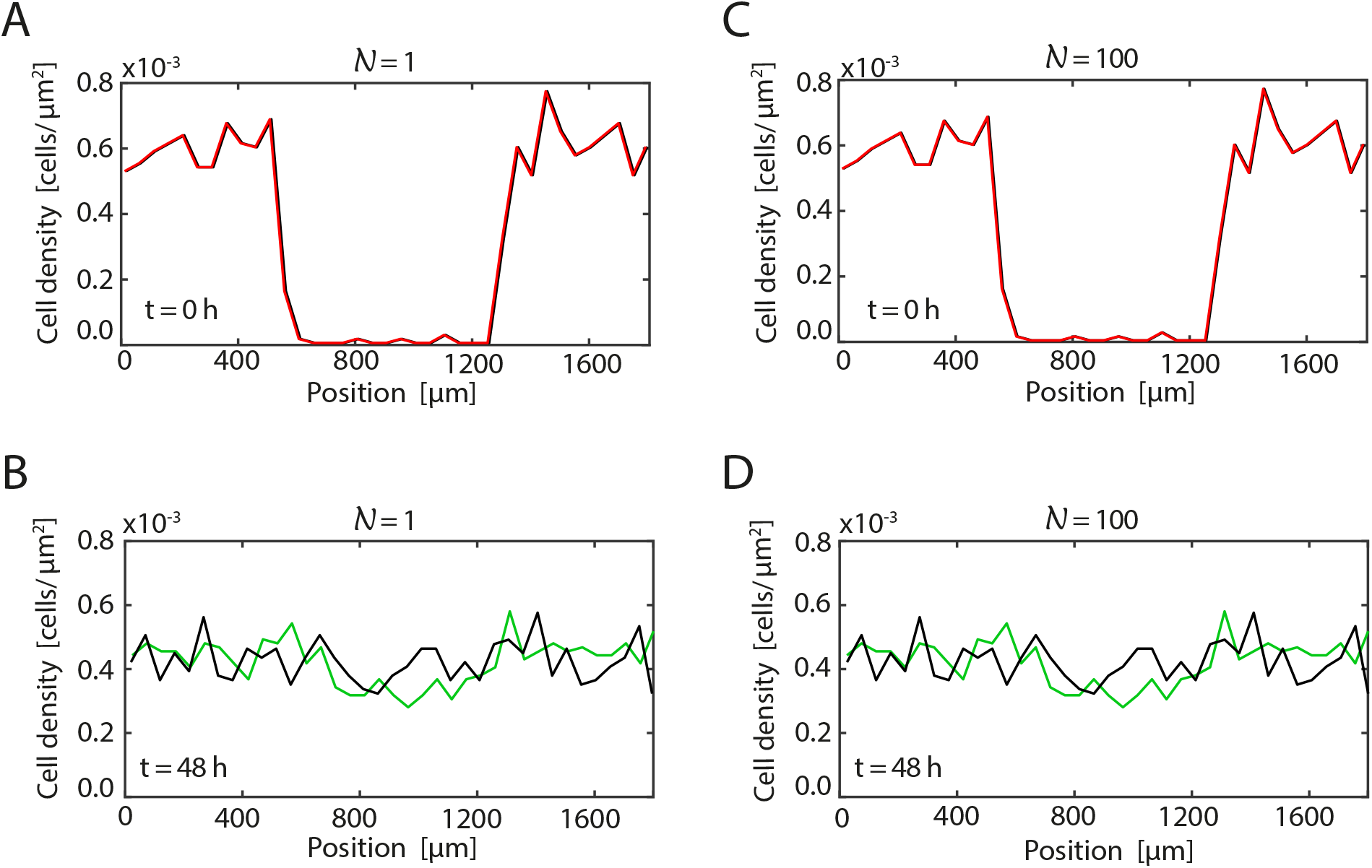
Comparison of the calibrated solutions of Model III and experimental profiles from experiment D. Results in A-B compare the experimental data (red, green) and Model II solution (black) at *t* = 0 and *t* = 48 h, respectively, for 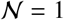 model realization. Results in C-D compare the experimental data (red, green) and Model II solution (black) at *t* = 0 and *t* = 48 h, respectively, averaged over 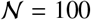 identically prepared realizations of the model. The best-fit parameter estimate is *f*_0_ = 0.08 *μ*m/h from Fig 13(A).

**Figure 15:**
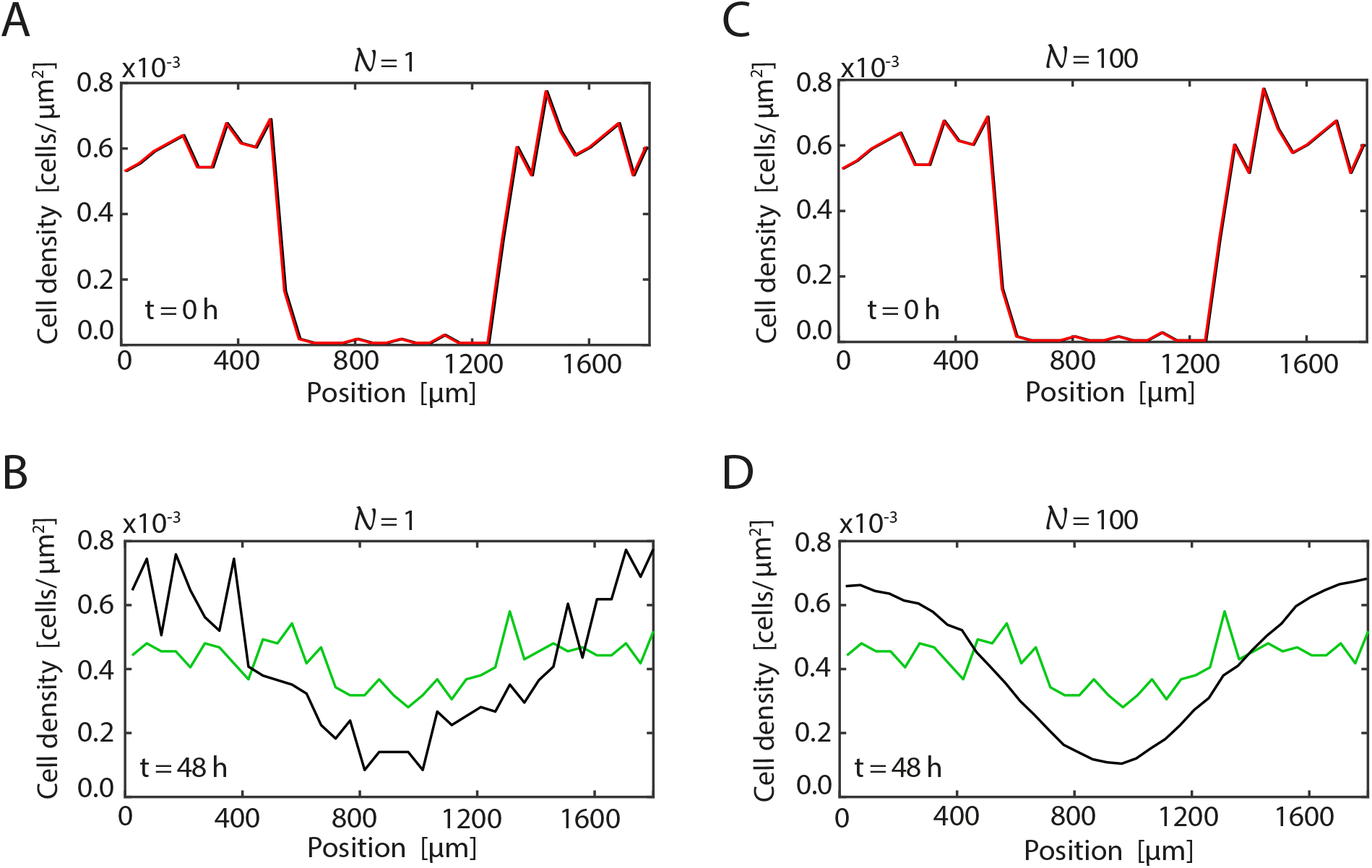
Comparison of the calibrated solutions of Model IV and experimental profiles from experiment D. Results in A-B compare the experimental data (red, green) and Model II solution (black) at *t* = 0 and *t* = 48 h, respectively, for 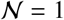 model realization. Results in C-D compare the experimental data (red, green) and Model II solution (black) at *t* = 0 and *t* = 48 h, respectively, averaged over 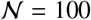 identically prepared realizations of the model. The best-fit parameter estimates is 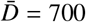 *μ*m^2^/h from Fig 13(B).

All results presented in this section of the main document focus on comparing the quality of match between Models I–IV and the experimental data using experimental replicate D. Similar comparisons between the best-fit solution of Models I–IV and experimental data from experimental replicates A, B and C are given in the Supporting Information (S3-S14 Fig). These additional comparisons are consistent with the comparisons made here in the main document.

## Conclusion

In this work we use a combined experimental-mathematical modelling approach to quantitatively explore the contribution of cell-to-cell interactions and random motility in driving the movement of cell fronts. We perform a series of IncuCyte ZOOM™ scratch assay experiments in which cells are pretreated with the chemotherapy drug, Mitomycin-C. This approach is useful because Mitomycin-C suppresses proliferation, thereby allowing us to focus on the role of cell migration in the experiments.

We quantitatively assess the role of cell-to-cell interactions, including short range pushing and longer range adhesion, by calibrating an off-lattice discrete stochastic model to match our experimental data set. The mathematical model that we use accounts for random cell motility, long range cell-to-cell attraction (adhesion), short range cell-to-cell repulsion (pushing) and dynamic cell size changes. We refer to this model as *Model I*. To explore the significance of these various cell-level mechanisms we systematically repeat the model calibration process for a range of simpler, more commonly used models. These simplified models account for: (i) random cell motility, long range cell-to-cell attraction (adhesion) and short range cell-to-cell repulsion (pushing) without any dynamical changes in cell size (*Model II*); (ii) long range cell-to-cell attraction (adhesion) and short range cell-to-cell repulsion (pushing) (*Model III*); and (iii) random cell motility only (*Model IV*).

Comparing the calibration of these four models to our experimental data provides insight into which model provides the most faithful representation of the experimental observation. Comparing estimates of 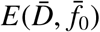 between the four models shows that Model I provides the best match to the experimental data. This result suggests that properly accounting for random motility, intercellular forces, including short range repulsion and longer range attraction, as well as dynamic changes in cell size, are important for this fairly typical experimental protocol. In contrast, calibrating Models II-IV to the data always provides estimates of model parameters that give the best match to the experimental data, but this does not mean that these simpler, more commonly used modelling frameworks, are the best model of the underlying biological processes. This is an important result because often in the mathematical biology literature a single type of model will be used to mimic an experiment, without investigating the more important question of whether that model provides a reasonable description of the underlying biological processes. Here, by systematically comparing the performance of four different, but related, mathematical models to our novel experimental data set, we provide insight into the underlying biological mechanisms in a way that is not possible when working with a single mathematical model in isolation. This approach of calibrating multiple competing mathematical models to match a single data set is a useful way to provide insight into the underlying biological processes, as well as providing insight into the important question of model selection [56, 57].

There are many ways that our study could be extended to provide additional insight. A key simplifying assumption that we make in our modelling of the experiments is that we give all agents in the simulations the same size at the beginning of the simulation, *δ*(0) = *δ*_0_. While our estimates of this initial size are based on experimental estimates given in Fig 2(C), our approach is to represent the distribution of observed cell sizes using a sample mean. A close examination of the experimental images in Fig 5 shows that there is considerable variability in the distribution of cell sizes at the beginning of the experiment. This variability is captured in the error bars in Fig 2(C) but neglected in our analysis. Therefore, a reasonable extension of the current analysis would be to incorporate this initial variability into the stochastic models with a view to understanding how this initial variability influences the population-level motion of the cell fronts. We leave this extension for future consideration.

